# Placental uptake and metabolism of 25(OH)Vitamin D determines its activity within the fetoplacental unit

**DOI:** 10.1101/2021.03.01.431439

**Authors:** Brogan Ashley, Claire Simner, Antigoni Manousopoulou, Carl Jenkinson, Felicity Hey, Jennifer M Frost, Faisal I Rezwan, Cory H White, Emma Lofthouse, Emily Hyde, Laura Cooke, Sheila Barton, Pamela Mahon, Elizabeth M Curtis, Rebecca J Moon, Sarah R Crozier, Hazel M Inskip, Keith M Godfrey, John W Holloway, Cyrus Cooper, Kerry S Jones, Rohan M Lewis, Martin Hewison, Spiros D Garbis, Miguel R Branco, Nicholas C Harvey, Jane K Cleal

**Author notes:** Corresponding author Dr Jane K Cleal; **Email:**. equal contributions (Brogan Ashley, Claire Simner). equal contributions (Nicholas C Harvey, Jane K Cleal). **Author Contributions:** JC, NH, MB, RL, CC: participation in study design execution and analysis. BA, CS, EL, EH, LC, RL, JC: participation in study experimental execution and analysis. CJ, FH, KJ, MH: Metabolism experimental design, analysis and interpretation. FR, CW, JH: Transcriptomic and DNA methylation analysis and interpretation. SB, PM, SRC, EC, RM, HI, KG, CC, NH: SWS data collection, analysis and interpretation. AM and SDG: Proteomics Experimental Design, Performed Proteomics analysis and data interpretation. JF: performed ChIP-seq experiments. MB: performed bioinformatics analyses. All: participation in manuscript drafting and critical discussion. **Data availability statement**: RNA sequencing data, ChIP-seq data and methylation array data that support the findings of this study have been deposited in the Gene Expression Omnibus (GEO) with the accession code GSE167431. All protein mass spectrometry data have been deposited to the ProteomeXchange Consortium via the Proteomics Identifications Archive with the data set identifier PXD011443.

## Abstract

Pregnancy 25-hydroxyvitamin D (25(OH)D) concentrations are associated with maternal and fetal health outcomes. Using physiological human placental perfusion and villous explants, we investigate the role of the placenta in regulating the relationships between maternal 25(OH)D and fetal physiology. We demonstrate active placental uptake of 25(OH)D3 by endocytosis, placental metabolism of 25(OH)D3 into 24,25-dihydroxyvitamin D3 and active 1,25-dihydroxyvitamin D [1,25(OH)2D3], with subsequent release of these metabolites into both the maternal and fetal circulations. Active placental transport of 25(OH)D3 and synthesis of 1,25(OH)2D3 demonstrate that fetal supply is dependent on placental function rather than simply the availability of maternal 25(OH)D3. We demonstrate that 25(OH)D3 exposure induces rapid effects on the placental transcriptome and proteome. These map to multiple pathways central to placental function and thereby fetal development, independent of vitamin D transfer. Our data suggest that the underlying epigenetic landscape helps dictate the transcriptional response to vitamin D treatment. This is the first quantitative study demonstrating vitamin D transfer and metabolism by the human placenta, with widespread effects on the placenta itself. These data demonstrate a complex interplay between vitamin D and the placenta and will inform future interventions using vitamin D to support fetal development and maternal adaptations to pregnancy.

## Introduction

Vitamin D (calciferol) cannot be synthesised by the fetus, so maternal vitamin D or its biologically significant metabolites 25-hydroxyvitamin D [25(OH)D] and/or 1,25-dihydroxyvitamin D [1,25(OH)_2_D] must be transferred across the placenta. This transfer is important for both fetal and lifelong health. Indeed maternal 25(OH)D concentrations are positively associated with fetal bone growth and birth weight [1, 2], and associations with bone health and body composition persist into postnatal life [3–7]. Therefore, understanding the regulatory mechanisms and rate-limiting steps of placental vitamin D transfer is a prerequisite for identifying options for targeted intervention to improve pregnancy outcomes. It is unclear whether the benefits of vitamin D and its metabolites are due to direct transfer to the fetus or indirect effects on the placenta. Fundamental questions need to be answered: **1)** does placental metabolism generate substantial quantities of vitamin D metabolites that contribute to the maternal or fetal circulation? **2)** How are these vitamin D metabolites taken up by placenta? And **3)** do vitamin D metabolites affect the placenta itself? This third question is particularly important as changes in placental function could subsequently influence the fetus independently of vitamin D transfer.

Maternal 25(OH)D was previously thought to diffuse passively across the placenta, and to be hydroxylated in the fetus to 1,25(OH)_2_D, as fetal metabolite levels relate to maternal levels for 25(OH)D [8–10] but not for 1,25(OH)_2_D [11, 12]. We challenge this idea, as work in the kidney questions the role of passive diffusion as a mechanism of vitamin D uptake and demonstrates that renal vitamin D uptake is driven primarily by receptor-mediated endocytosis of vitamin D bound to vitamin D binding protein (DBP) [13] or albumin [14, 15]. We propose a similar endocytic uptake mechanism for vitamin D (25(OH)D and 1,25(OH)_2_D) in the placenta, indicating an active role for the placenta in regulating fetal 25(OH)D delivery. Furthermore, placental metabolism of maternal 25(OH)D, either by placental 1α-hydroxylase (*CYP27B1*) metabolism into 1,25(OH)_2_D or 24-hydroxylase (*CYP24A1*)-mediated breakdown, will influence the quantity and form of vitamin D metabolite reaching the fetus. In human placenta, both enzymes are localised to the syncytiotrophoblast, which is the primary barrier to maternal-fetal exchange. Whether the activity of these placental enzymes generates substantial quantities of vitamin D metabolites that significantly contribute to the maternal or fetal circulations is unknown.

Placental uptake and metabolism of 25(OH)D and 1,25(OH)_2_D will determine the availability of 1,25(OH)_2_D for inducing transcriptional changes within the placenta via the vitamin D receptor (VDR), retinoid X receptor-alpha (RXRA) receptor dimer. Placental 1,25(OH)_2_D may therefore regulate gene pathways involved in placental functions that influence fetal growth and risk of lifelong poor health. Indeed, we have shown that maternal 25(OH)D and DBP concentrations relate to expression of placental genes, including those associated with fetal and postnatal lean mass [16, 17]. The effects of vitamin D, thought to be via the 1,25(OH)_2_D form, on placental gene expression are not clear as these are cell/tissue specific and depend on the chromatin state and DNA-binding proteins available [18, 19]. Furthermore, vitamin D may induce epigenetic regulation of gene expression. For example, in immune cells ligand-bound VDR exerts epigenetic effects via interaction with chromatin modifiers and coactivator or corepressor proteins, such as histone modifiers [18, 19]. Changes to DNA methylation have also been reported with vitamin D cellular incubation, patient supplementation [19–21], or in association with vitamin D status [22], supporting a role of VDR in altering DNA methylation.

The current study was designed to establish how 25(OH)D_3_ is taken up, metabolised and mediates gene expression within the human placenta using the more physiological perfusion model and intact villous explant culture systems, contrasting with the cell model approach of previous studies. We show novel active mechanisms of 25(OH)D_3_ uptake by the human placenta and placental metabolism of 25(OH)D_3_ influencing fetal and maternal levels. Furthermore, 25(OH)D induces placental specific effects on the transcriptome, the native and *in vivo* modified proteome expressed in patterns relevant to placental function and therefore fetal development, independent of vitamin D transfer. These effects are dependent on the underlying epigenetic landscape. By discovering the mechanisms underlying the complex interactions between maternal vitamin D levels, placental vitamin D handling and genetic/epigenetic processes in the placenta, our findings inform the identification of functional biomarkers that relate to fetal growth and the risk of disease across the life course.

## Results and Discussion

### 25(OH)D_3_ is taken up by the human placenta and both metabolised and transferred into the fetal circulation

The use of ^13^C-labelled 25(OH)D_3_ in the maternal circulation of the *ex vivo* placental perfusion model allowed actual quantification of the placental metabolism of ^13^C-25(OH)D_3_ and transfer of this ^13^C-25(OH)D and its metabolites from the maternal to fetal circulation in the term placenta. Although *in vivo* maternal and fetal circulating 25(OH)D_3_ levels at term may correlate [23], the level of actual 25(OH)D transfer and metabolism by the human placenta is unclear as prior data provided inconsistent results using *in vivo* plasma measures or closed loop perfusion experiments [24]. We investigated the metabolism of ^13^C-25(OH)D_3_ within placental tissue and transfer across the placenta using the whole placental cotyledon perfusion open loop methodology as adapted in our laboratory [17]. Using five fresh placentas, ^13^C-25(OH)D_3_ (plus albumin as a binding protein) was perfused for 5 h into the maternal circulation of a single isolated cotyledon and the amount transferred into the placenta and fetal circulation quantified using liquid chromatography hyphenated with mass spectrometry (LC-MS/MS). The amount of each specific vitamin D metabolite transferred into the fetal and maternal circulation was also quantified using LC-MS/MS or Enzyme Immunoassay (EIA).

Following 5 h of placental perfusion there was loss of maternal ^13^C-25(OH)D_3_ from the maternal circulation indicative of placental uptake (27.4 ± 6.3% of maternal stock) and release into the fetal circulation (5.1 ± 1.5% of maternal stock; **Figure 1a**). These amounts are consistent with the levels taken up by the kidney via active mechanisms [25]. The transfer rate of ^13^C-25(OH)D_3_ into the placenta was 3.01 ± 1.12 pmol/min/g, with 0.45 ± 0.27 pmol/min/g stored in placental tissue, 2.25 ± 0.97 pmol/min/g metabolised and 0.30 ± 0.10 pmol/min/g transferred to the fetal circulation (**Figure 1b**). However, there was no correlation between ^13^C-25(OH)D_3_ levels in the maternal and fetal circulations of this open loop placental perfusion system: a system that identifies delivery, but not build-up, of substrates in either circulation (**Figure 1g**). Together these data suggest that the placenta plays an important role in determining the amount of 25(OH)D_3_ transferred to the fetus and also stores and uses 25(OH)D_3_ for its own cellular needs. Placental metabolism and uptake of ^13^C-25(OH)D_3_ are therefore key factors in determining fetal ^13^C-25(OH)D_3_ supply (**Figure 2a**).

**Figure 1.**
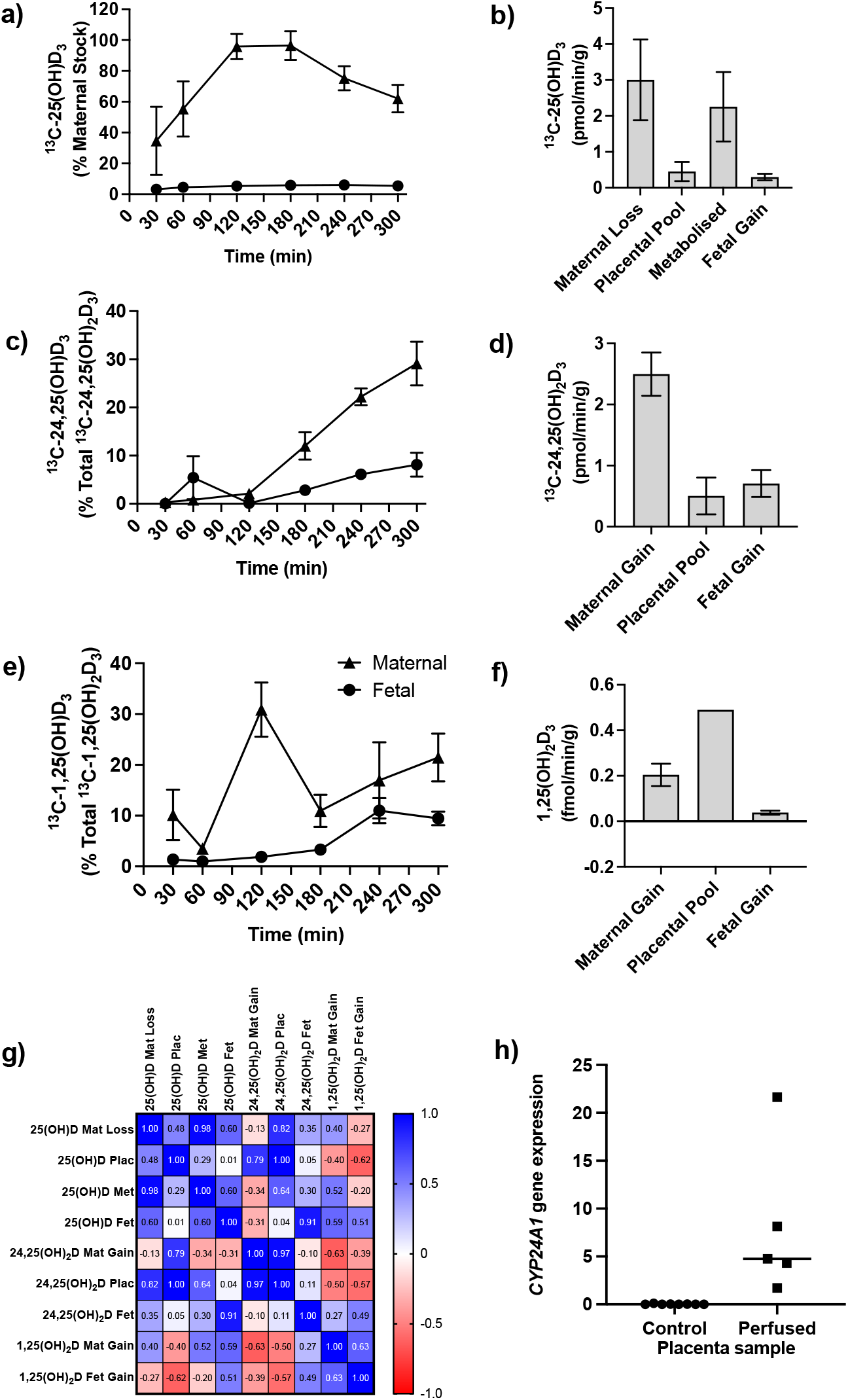
Transfer and metabolism of ^13^C-25(OH)D_3_ by the term perfused human placenta over 5 h. **a)** ^13^C-25(OH)D_3_ in maternal and fetal circulations as a % of maternal perfusate concentration. **b)** Rate of maternal ^13^C-25(OH)D_3_ lost from the maternal circulation, accumulated in placental tissue, metabolised or transferred to the fetal circulation. **c)** ^13^C-24,25(OH)_2_D_3_ transfer into the maternal and fetal circulations as a % of ^13^C-24,25(OH)_2_D_3_ metabolised by the placenta. **d)** Rate of placental production of ^13^C-25(OH)D_3_ and transfer rate into the maternal and fetal circulations. **e)** 1,25(OH)_2_D_3_ transfer into the maternal and fetal circulations as a % of 1,25(OH)_2_D_3_ produced by the placenta. **f)** Rate of placental production of ^13^C-1,25(OH)_2_D_3_ and 1,25(OH)_2_D_3_ transfer into the maternal and fetal circulations. **g)** Pearson’s correlations between metabolites. **h)** *CYP24A1* relative mRNA expression in the 25(OH)D perfused placental samples compared to nonperfused control placental tissue samples. Data are presented as mean (SEM).

**Figure 2:**
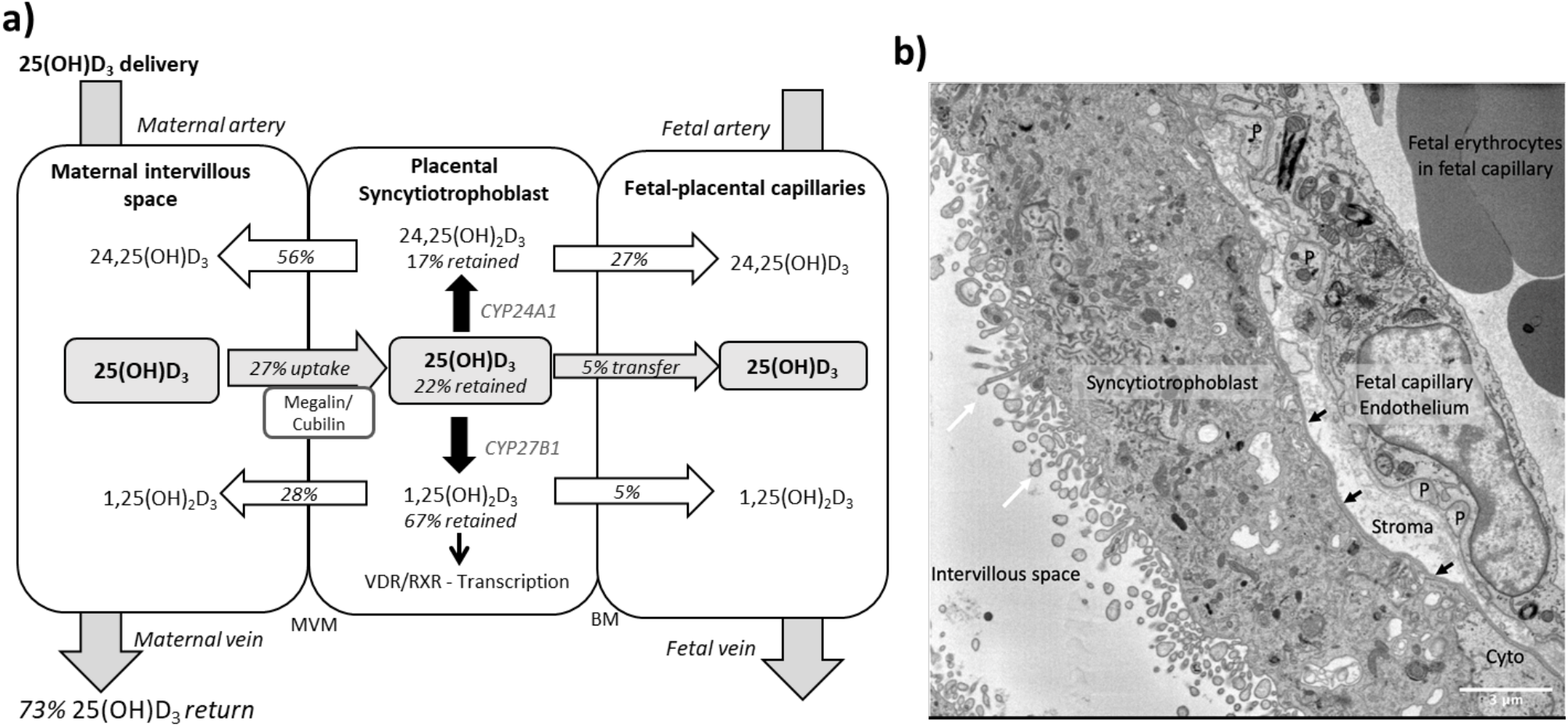
Summary of vitamin D transport and metabolism by the placenta. **a)** Schematic model of ^13^C-25(OH)D_3_ transfer and metabolism by the term perfused human placenta over 5 h. **b)** Electron microscopy image showing a cross section of the human placental barrier at term. The intervillous space is filled with maternal blood, the syncytiotrophoblast forms a continuous barrier across the surface of the villi, the microvilli on the apical plasma membrane are indicated by white arrows and the syncytiotrophoblast basal membrane can be seen abutting the trophoblast basal lamina which is indicated by black arrows. A small region of cytotrophoblast can be seen labelled ‘cyto’ between the syncytiotrophoblast and trophoblast basal lamina. The connective tissue of the villous stroma lies between the trophoblast and the fetal capillaries. The stroma also contains fibroblasts and macrophages which are not shown here. Pericyte fingers around the fetal capillary are labelled ‘P’. The fetal capillary endothelial cells form the fetal blood vessel.

The primary barrier to transfer, the placental syncytiotrophoblast, expresses the vitamin D-metabolising enzymes 24-hydroxylase (*CYP24A1*) and 1α-hydroxylase (*CYP27B1*) [26]. There was direct evidence of 24-hydroxylase action in the placental tissue with metabolism of 25(OH)D_3_ into ^13^C-24,25(OH)_2_D_3_ (**Figure 1c & 1d**) but no detectable epi25(OH)D_3_. Placental ^13^C-25(OH)D_3_ and ^13^C-24,25(OH)_2_D levels were positively correlated (r = 0.998, p = 0.0001; **Figure 1g**). Of this ^13^C-24,25(OH)_2_D_3_, 16.78 ± 7.20% (0.50 ± 0.30 pmol/min/g) was stored in the placental tissue, 20.4 ± 3.21% (0.71 ± 0.10 pmol/min/g) was transferred into the fetal circulation and 66.56 ± 6.31% (2.05 ± 0.35 pmol/min/g) was transferred into the maternal circulation (**Figure 1c & 1d**). There was also evidence of placental 1α-hydroxylase activity with metabolism of 25(OH)D_3_ into 1,25(OH)_2_D_3_ (**Figure 1e & f**). Transfer of 1,25(OH)_2_D_3_ into both the maternal (0.20 ± 0.05 fmol/min/g) and fetal (0.04 ± 0.01 fmol/min/g) circulations occurred over the 5 h experiment with direct evidence of placental metabolism of ^13^C-25(OH)D_3_ into ^13^C-1,25(OH)_2_D_3_ (0.49 fmol/min/g, n = 1 due to detection limits). These data provide evidence that once within the placenta 25(OH)D can be metabolised into downstream products that could have effects within the placenta itself including VDR-mediated transcription.

We observed greater release of ^13^C-25(OH)D_3_ metabolites into the maternal circulation compared with the fetal circulation (**Figure 2a**). The transfer of ^13^C-25(OH)D_3_ and its metabolites in the placenta to fetal direction may be more challenging, and in particular, diffusion across the water-filled villous stroma may prove a barrier to their diffusion (**Figure 2b**); consistent with recent observations on placental cortisol transfer [27]. Indeed, positive correlations between fetal ^13^C-25(OH)D_3_ and ^13^C-24,25(OH)_2_D concentrations (r = 0.913, p = 0.03 **Figure 1g**) suggest similar or limited transfer processes for these metabolites in the placental to fetal direction. This is supported by the facts that placental to maternal transfer rates were higher, prioritising maternal supply, and that placental and maternal ^13^C-24,25(OH)_2_D levels positively correlated (r = 0.966, p = 0.03; **Figure 1g**). The greater amount of ^13^C-25(OH)D_3_ metabolites released into the maternal circulation, compared with the fetal circulation, also indicates a potential role of the placenta in vitamin D-mediated maternal adaptations to pregnancy. These data substantiate a role for placental metabolism and transfer mechanisms being important for fetal vitamin D metabolite levels. These experiments therefore show that the placenta can produce substantial quantities of 25(OH)D_3_ metabolites, which can contribute to both the maternal and fetal circulating levels and highlights the importance of placental expression of the vitamin D-metabolising enzymes 24-hydroxylase (gene *CYP24A1*) and 1α-hydroxylase (gene *CYP27B1*).

Placental *CYP24A1* gene expression (which is induced by 1,25(OH)_2_D_3_) was significantly increased following perfusion with ^13^C-25(OH)D_3_ compared with control (**Figure 1h**), providing further evidence that 25(OH)D_3_ is converted to active 1,25(OH)_2_D_3_ by 1α-hydroxylase in human placenta. In a subset of term human placentas taken from the Southampton Women’s Survey (SWS, n = 102), we did not observe an association between maternal 34-week pregnancy plasma 25(OH)D_3_ levels and *CYP24A1* (r = −0.15, p = 0.18) or *CYP27B1* (r = 0.14, p = 0.20) gene expression [16, 28]. In the SWS placentas, *CYP24A1* mRNA expression was associated with maternal mid-upper arm muscle area pre-pregnancy (r = 0.33, p = 0.001 n = 101), at 11 weeks’ gestation (r = 0.33, p = 0.02, n = 76) and at 34 weeks’ gestation (r = 0.26, p = 0.01, n = 95). This suggests that factors other than vitamin D levels, such as maternal body composition, may regulate baseline levels of this gene. The fact that the placenta can produce 1,25(OH)_2_D_3_, which induces expression of the degradation enzyme *CYP24A1* suggests homeostatic regulation of placental vitamin D activity and potentially its transfer to the fetus. This has implications for the effectiveness of vitamin D supplementation during pregnancy.

### Uptake of 25(OH)D_3_ and albumin into the placenta is prevented by inhibition of endocytosis

To explore the mechanisms of placental 25(OH)D_3_ uptake, fresh term human placental villous fragments were incubated for 8 h with 20 μM 25(OH)D_3_ plus the binding protein albumin. Analysis by qPCR revealed a significant increase in placental *CYP24A1* gene expression in villous fragments following incubation with 25(OH)D_3_ compared to control (**Figure 3a**). In the presence of albumin, *CYP24A1* mRNA expression was increased compared to 25(OH)D_3_ alone, suggesting that albumin may facilitate 25(OH)D_3_ uptake (**Figure 3a**). We suggest that receptor-mediated endocytosis may play an important role in the uptake of vitamin D into the human placenta as seen in the kidney. The increased *CYP24A1* expression with vitamin D uptake is increased further by the presence of a binding protein; this could be due to increased solubility or receptor-mediated binding for the uptake mechanism.

**Figure 3.**
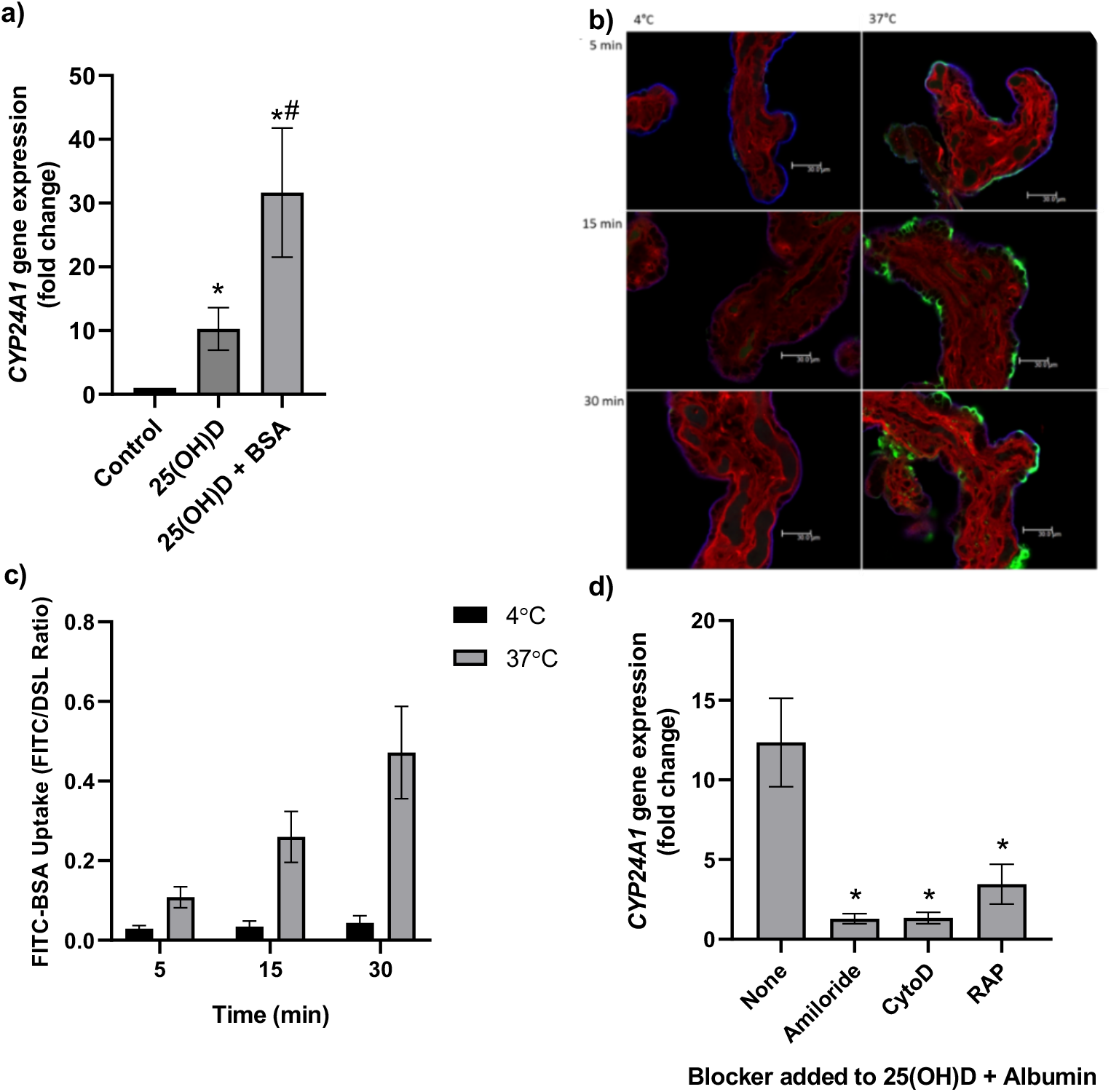
Uptake of 25(OH)D_3_ in placental villous fragments is facilitated by albumin and mediated by endocytic processes. **(a)** Relative *CYP24A1* mRNA expression was increased in placental villous fragments incubated with 25(OH)D_3_ (n = 15) compared to control (ethanol plus albumin, n = 15; *p < 0.001), and was increased further with 25(OH)D_3_ and albumin (n = 6) compared to 25(OH)D_3_ alone (^#^p < 0.05). Uptake of FITC-albumin to placental fragments is mediated by endocytic mechanisms. **(b)** Representative images showing FITC-albumin uptake into placental villous fragments at 5, 15 and 30 min. Green, FITC-albumin; red, villous stroma stained by rhodamine-PSA; and blue, MVM stained by biotin-DSL. **(c)** FITC-albumin uptake increased with time (p = 0.03) and temperature (p = 0.03), n = 3. **(d)** *CYP24A1* gene expression was reduced by addition of the blockers amiloride, cytochalasin D (CytoD) and Receptor Associated Protein (RAP) compared to 25(OH)D plus albumin-stimulated expression with no blocker (* p < 0.05; n = 4-5). BSA, bovine serum albumin. Data are presented as mean (SEM).

The placental uptake mechanisms for 25(OH)D_3_ via the carrier protein albumin were investigated by measuring uptake of FITC-labelled albumin into placental villous fragments. Uptake over 30 min at 4°C (a temperature that blocks endocytosis) and 37°C showed a significant effect of both time and temperature on placental FITC-albumin uptake, indicating an active uptake mechanism (**Figures 3b and 3c**). In addition, FITC-dextran uptake was not observed suggesting the albumin uptake mechanism is selective (data not shown).

We next investigated whether binding protein-mediated vitamin D uptake occurred via endocytosis. In the kidney, both DBP and albumin bind to the plasma membrane receptors megalin (LDL-related protein 2 (*LRP2*)) and cubilin (*CUBN*) [29, 30], which mediate vitamin D internalisation by clathrin-dependant endocytosis [25, 31]. Both receptors are expressed in the primary barrier to transfer, the syncytiotrophoblast, suggesting a role for vitamin D uptake by the placenta [26]. To investigate this, fresh placental villous fragments (n = 4-5 placentas per treatment) were incubated with 20 μM 25(OH)D_3_ and albumin, plus the endocytic inhibitors 5 mM amiloride, 10 μM cytochalasin D and 2 μM Receptor Associated Protein (RAP) which block pinocytosis, classical clathrin-dependent endocytosis and megalin-mediated endocytosis, respectively. The vitamin D-mediated up-regulation of *CYP24A1* mRNA expression was significantly reduced by amiloride, cytochalasin D and RAP in the presence of albumin (**Figure 3d**). These data suggest that 25(OH)D_3_ is taken up by the human placenta via a binding protein using an endocytic mechanism that is blocked by RAP, thus indicating a role via megalin. This suggests that vitamin D is transported into the placenta by a selective and controlled active mechanism, not by simple diffusion as previously thought, which has potential implications for the effectiveness of maternal vitamin D supplementation and its bioavailability to the fetus.

The gene expression of megalin/*LRP2* and cubilin/*CUBN* were investigated in term human placentas from the SWS [28]. Both showed no association with maternal 34-week pregnancy plasma 25(OH)D_3_ levels (data not shown) and positive associations with measures of fetal size; *LRP2* mRNA expression related to crown-rump length at 11 weeks’ gestation (r = 0.23, p = 0.05, n = 51) and birth weight (r = 0.20, p = 0.05, n = 102); *CUBN* mRNA expression related to birth head circumference (r = 0.22, p = 0.03, n = 102) and crown-heel length (r = 0.21, p = 0.04, n = 102). This supports a role for the placenta in mediating the amount of vitamin D transferred to the fetus, which will be determined by placental uptake capacity, which in turn effects fetal size. We suggest that the placenta may be playing an active role in mediating the supply of vitamin D to the fetus thus contributing to its regulation and effects on development.

Vitamin D induced gene expression changes in the placenta could also mediate fetal development by altering placental function. Placental uptake and metabolism of 25(OH)D_3_ and 1,25(OH)_2_D_3_ will ultimately determine the amount of 1,25(OH)_2_D_3_ available within the placenta for activation of the VDR and initiation of transcription. Interestingly, in SWS placentas we see associations between VDR mRNA expression and measures of fetal size suggesting that vitamin D-mediated transcription is important for regulating aspects of placental function that affect fetal size. Placental *VDR* mRNA levels associate negatively with measures of fetal and neonatal size and body composition (**Table 1**), and *RXRA* mRNA levels associate negatively with 4-year lean mass (r = −0.27, p = 0.005) and fat mass (r = −0.34, p = 0.03). The placental levels of VDR and RXR may therefore determine the utilisation of available 1,25(OH)_2_D by the placenta via VDR binding. Increased VDR/RXR levels will result in activation of the *CYP24A1* gene encoding the enzyme that breaks down 25(OH)D and 1,25(OH)_2_D, reducing the quantities of the two metabolites available for transfer to the fetus for mediating effects on growth.

**Table 1.**
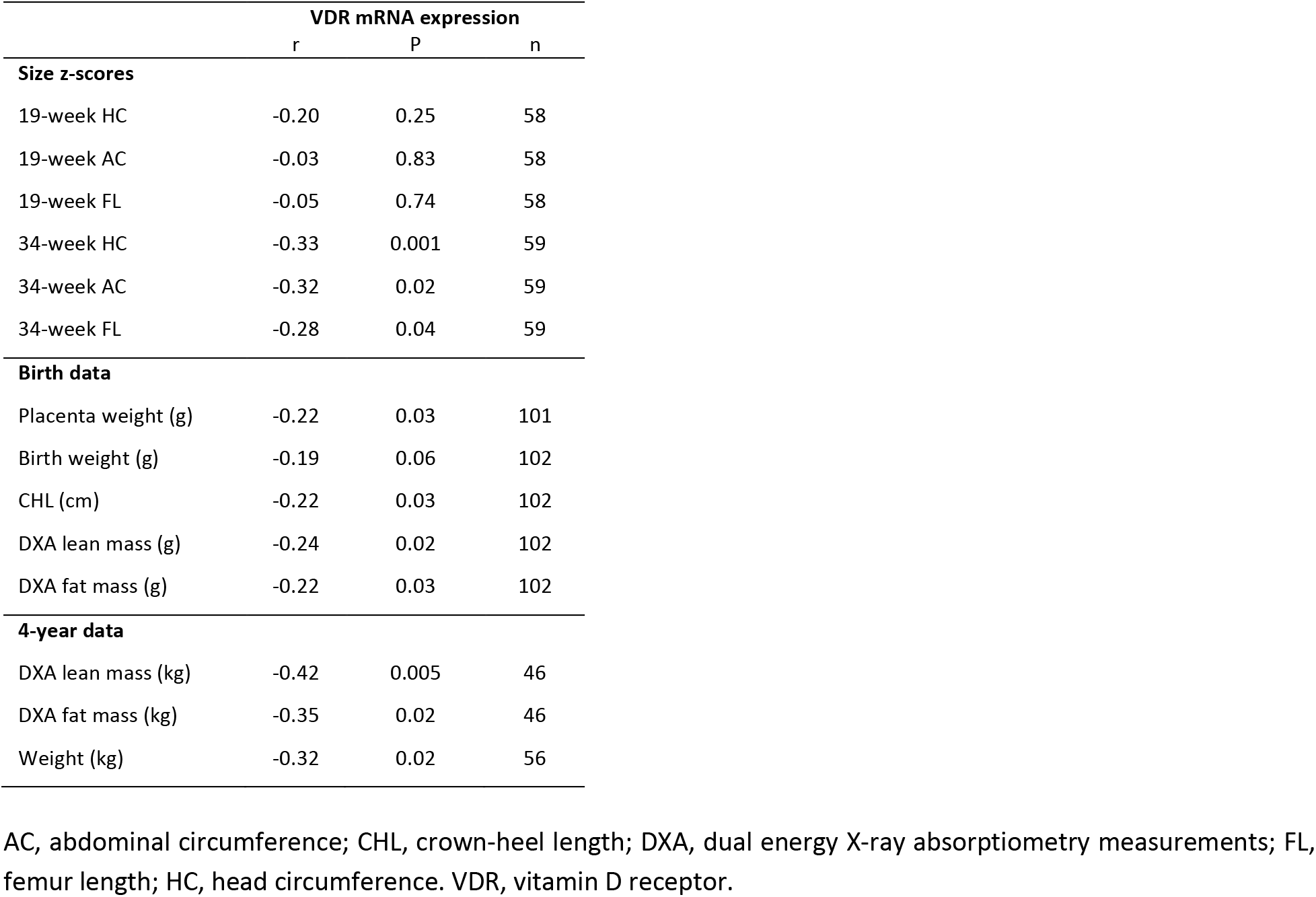
Associations of offspring size and vitamin D receptor relative mRNA expression in the Southampton Women’s Survey.

### Short-term 25(OH)D_3_ exposure leads to transcriptomic changes in human placenta

Our previous experiments show that conversion of 25(OH)D_3_ into 1,25(OH)_2_D_3_ within the human placenta initiates gene transcription of a specific vitamin D-responsive gene, *CYP24A1*. To understand the genome-wide effects of vitamin D on the placenta, RNA sequencing analysis was performed on villous fragments from four fresh term human placentas incubated with 20 μM 25(OH)D_3_ plus albumin for 8 h compared to control.

Principal component analysis (PCA) demonstrated condition clustering indicating an effect of treatment (**Figure 4a**). Placental fragments exposed to 25(OH)D_3_ displayed marked changes in gene expression compared to those in control buffer (**Figure 4b**). Using a 5% false discovery rate (FDR) we observed 493 genes to be differentially expressed between 25(OH)D_3_ treated compared to control placental fragments (358 increased and 135 decreased; **Supplementary Table 1**) and those with a fold change of 2 or more are presented (**Figure 4c**). We compared the gene expression changes observed following short term vitamin D exposure with those in a published dataset from human placenta (GSE41331), which looked at longer-term vitamin D response (24 h). Whilst considerable differences were expected and observed, it was reassuring that genes found to be upregulated at 8 h also tended to have increased expression in the 24 h dataset, whereas downregulated genes were unchanged (**Supplementary Figure 1**). This probably reflects the fact that upregulated genes are the likely direct targets of VDR. Differentially expressed genes in common between both placental datasets included *CYP24A1* and *SNX31* (**Supplementary Figure 1**).

**Figure 4:**
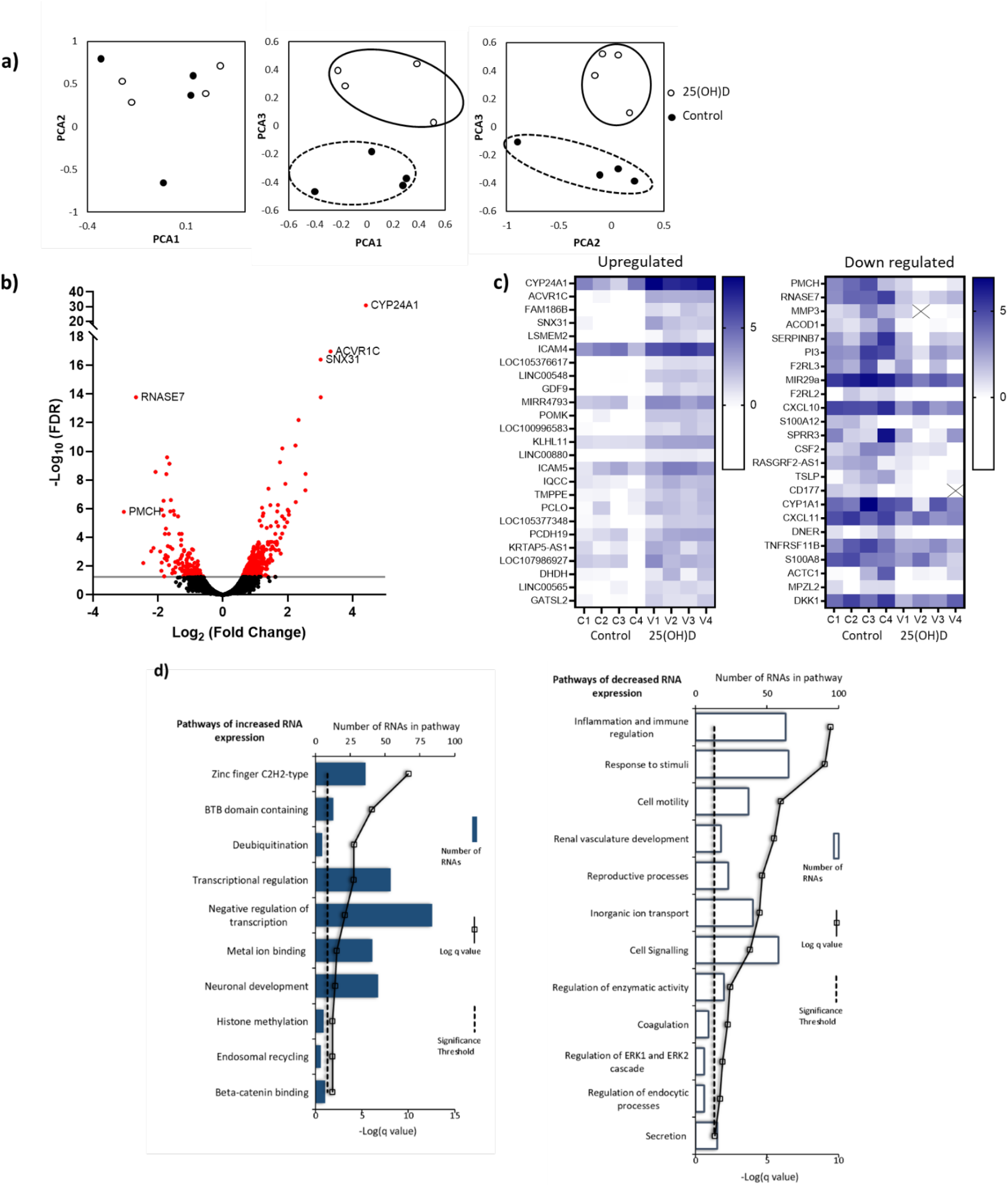
RNA-seq derived gene signatures of human placental samples following 8 h 25(OH)D_3_ incubation. **a)** Principal component analysis indicated clustering of 25(OH)D_3_ treated samples (clear) compared with control samples (black). **b)** Volcano plot showing 493 genes were differentially expressed (red) at a FDR adjusted p-value < 0.05 (grey line). **c)** Differentially expressed genes with a fold change of 1.5 or above at FDR 0.05 are presented as heatmaps (log2(normalised expression)). **d)** Pathway analysis (Toppgene) of all differentially expressed genes (no fold-change cut-off) reveals both up and down regulation of molecular function and biological process gene pathways following 8 h 25(OH)D_3_ incubation. For pathway analysis, significance was adjusted using the Benjamini-Hochberg correction depicted by −log(B&H q-value) with a significance threshold of 1% (dashed line).

To identify the biological functions regulated by vitamin D in human placenta we performed pathway analysis incorporating all differentially expressed genes using Toppgene with the ToppFun sub-program, significance was adjusted using the Benjamini-Hochberg correction depicted by −log(B-H p-value) and a significance threshold of 1%. The pathway analysis revealed that genes from the pathways involved in cytokine binding and immune or inflammatory responses were downregulated following 25(OH)D_3_ treatment (**Figure 4d**). These findings fit with the role of vitamin D in regulating cellular growth and immune function in a protective manner [18]. Genes from gene pathway terms related to transcriptional regulation were upregulated following 25(OH)D_3_ treatment (**Figure 4d**). These included many different transcription factors (e.g. *REST, FOXO3*), suggesting that this early response to 25(OH)D_3_ treatment maybe to cascade the signal to other transcriptional regulatory networks. Furthermore, transcription factors known to interact with VDR, such as the central regulators of gene expression CREB-binding protein (*CREBBP*) and its paralog *EP300*, were upregulated supporting the idea that the response to 25(OH)D_3_ treatment is about potentiating a robust secondary response. Effects on key transcription factors such as *CREBBP* and *EP300*, which can also interact with enhancer-bound transcription factors to activate transcription and have histone acetyltransferase activity, indicates that epigenetic mechanisms may play a role in 25(OH)D_3_ mediated regulation of gene expression.

### The underlying epigenetic landscape helps to dictate the transcriptional response upon vitamin D treatment

Short-term 25(OH)D_3_ exposure leads to transcriptomic changes in human placenta, however the role of epigenetic factors in this observation is unclear. We therefore investigated whether 8 h incubation with 20 μM 25(OH)D_3_ plus albumin altered DNA methylation in villous fragments from eight fresh term human placenta compared to control. DNA methylation was measured using the Illumina EPIC 850K array and CpGs with altered DNA methylation following 25(OH)D_3_ treatment were identified using a Wilcoxon signed-rank test (p < 0.05).

Short-term 25(OH)D_3_ exposure led to limited alterations in DNA methylation, with 319 CpGs displaying methylation differences larger than 10% (230 hypomethylated, 89 hypermethylated; **Supplementary Table 2**). Most of these changes involved isolated CpGs, whereas clusters of 2 or more differentially methylated CpGs were only observed in two hypomethylated and six hypermethylated regions (**Figure 5a**). Notably, the promoters of the upregulated genes identified in the RNA-seq data, displayed markedly lower methylation than those of downregulated genes, in both control and 25(OH)D_3_-treated conditions (**Figure 5b**). Consistent with this observation, 72% of upregulated gene promoters overlapped with CpG islands, in contrast with 23% of the downregulated (and 56% genome-wide). These results suggest that the underlying epigenetic landscape helps to dictate the transcriptional response to 25(OH)D_3_ treatment, possibly by enabling VDR binding at open chromatin regions.

**Figure 5:**
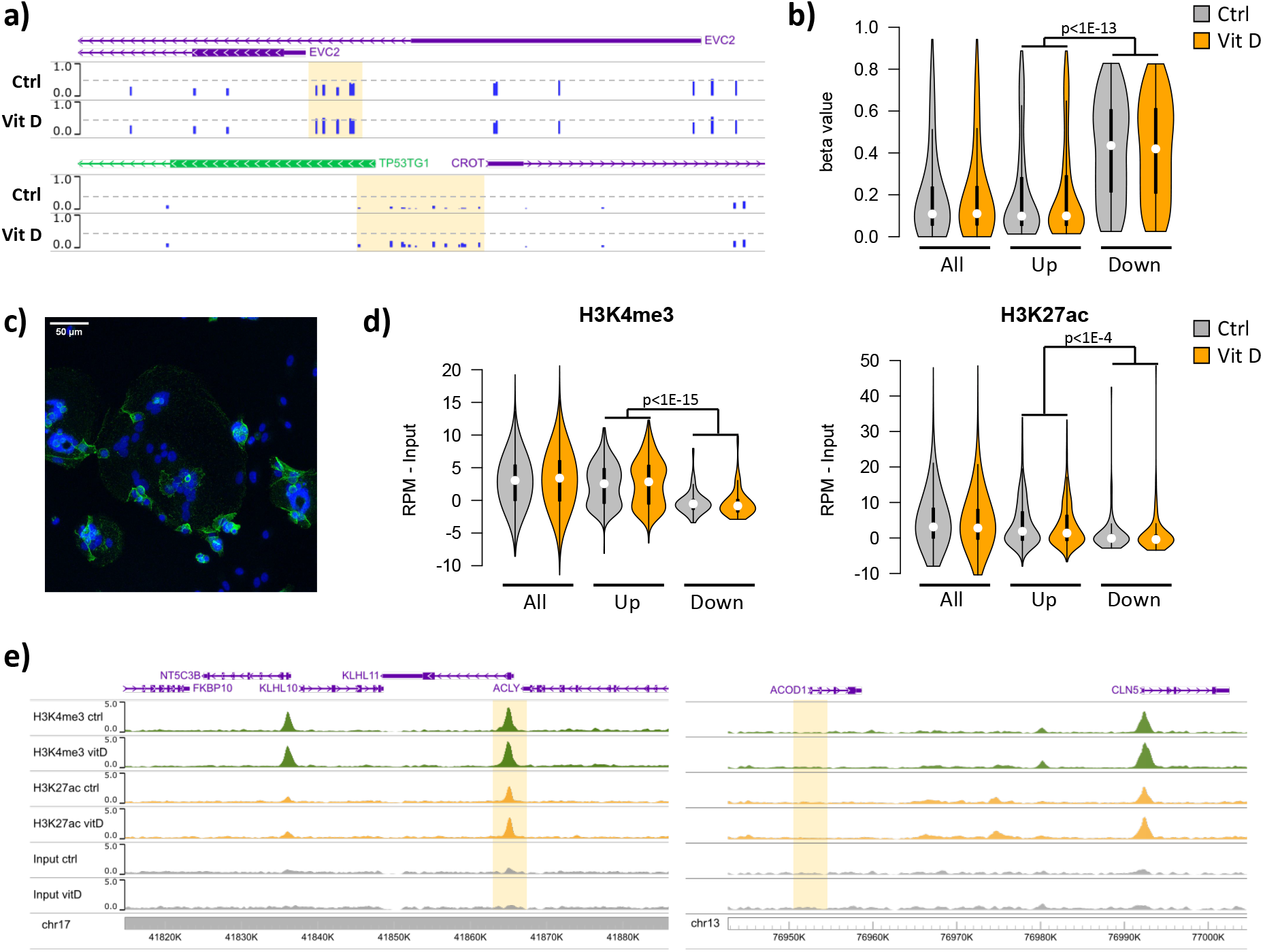
Short term vitamin D exposure has limited effects on placental methylation but the pre-existing epigenetic landscape has a major effect on vitamin D mediated transcription. **a)** Placental fragments were exposed for 8 h to 25(OH)D_3_, which led to limited alterations in DNA methylation. Shown are two examples of clusters of hypermethylated CpGs (highlighted in yellow), where the blue bars represent the array’s beta value for individual CpGs. **b)** The promoters of the upregulated genes identified in the RNA-seq data, displayed lower methylation than those of downregulated genes, in both control and 25(OH)D_3_ treated conditions. To extend these observations, we performed ChIP-seq on syncytialised cytotrophoblast cells incubated with 20 μM 25(OH)D_3_ for 24 h (n = 2 placentas). **c)** Representative confocal microscopy image of cytotrophoblast cells cultured for 90 h and stained with DAPI (blue; nuclei) and desmoplakin (green), present on the cell surface. Multiple nuclei within a single cell demonstrate syncytialisation has occurred. **d)** The promoters of upregulated genes (identified in the RNA-seq data) displayed higher levels of both H3K4me3 and H3K27ac than those seen at downregulated genes. **e)** Examples of specific upregulated (KLHL11) and downregulated (ACOD1) genes, showing no changes in the enrichment of H3K4me3 or H3K27ac at the promoter (highlighted in yellow) when comparing control and vitamin D conditions.

To extend these observations, we performed ChIP-seq on syncytialised cytotrophoblast cells [32] incubated with 20 μM 25(OH)D_3_ for 24 h (n = 2 placentas; **Figure 5c**). We profiled histone modifications associated with open chromatin, H3K4me3 and H3K27ac, the latter being also an *in vivo* target of EP300/CREBBP acetyltransferase activity. We detected no loci with significant changes in the enrichment of these marks when comparing control and vitamin D conditions (see methods for details). However, in line with the DNA methylation data, we found that promoters of upregulated genes displayed strikingly higher levels of both H3K4me3 and H3K27ac than those seen at downregulated genes (**Figure 5d**). This pattern was also seen when focusing only on CpG island promoters (**Supplementary Figure 2a**), ruling out the possibility that higher levels of H3K4me3/H3K27ac at upregulated genes were only due to increased CpG content, and thus reinforcing the idea that open chromatin poises genes for upregulation by VDR. Although our ChIP-seq data are from isolated cytotrophoblast, we observed very similar patterns in ENCODE data from 16-week placenta (**Supplementary Figure 2b**).

Given these observations, we asked whether a preference for open chromatin was specific to the VDR transcriptional response in placenta. We therefore analysed gene expression and open chromatin (FAIRE-seq) data from THP1 (leukemic monocyte) cells treated with 1,25(OH)_2_D_3_ for 4 h [33]. The transcriptional response in THP1 cells was dramatically different from that seen in the placenta, with only 5 upregulated and 8 downregulated genes in common. Interestingly, despite this large difference, the promoters of both placenta- and THP1-upregulated genes displayed open chromatin in THP1 cells (**Supplementary Figure 2c**), suggesting that other factors determine the specificity of the vitamin D response in these cells. In contrast, H3K4me3 levels in the placenta were higher for promoters of placenta-upregulated genes than for THP1-upregulated ones (**Supplementary Figure 2d**), suggesting that the 25(OH)D_3_ response in the placenta is partially guided by the chromatin status. Importantly, these results suggest that an altered epigenetic landscape in the placenta may lead to differences in the response to 25(OH)D_3_ exposure.

Our data suggest that in human placenta 25(OH)D_3_ is metabolised to the active 1,25(OH)_2_D_3_, which can regulate gene expression at several levels in an intracrine fashion within the same tissue. Whether the observed 25(OH)D_3_-induced gene expression changes followed through into protein expression within this short time frame or whether post-translational protein modifications could also be observed in 25(OH)D_3_ treated placental tissue was explored using LC-MS/MS.

### Short-term 25(OH)D_3_ exposure leads to proteomic changes in human placenta

The quantitative proteomic analysis of term placental villous fragments exposed for 8 h to 25(OH)D_3_ plus albumin (20 μM, n = 4 placentas) compared with patient-matched controls profiled 9279 unique protein groups (peptide level FDR, p < 0.05). Of these proteins, 246 were differentially expressed following 8 h exposure to 25(OH)D_3_ compared with control (paired permutation testing, p < 0.05). 25(OH)D_3_ incubation resulted in differential expression of 98 methylated, 67 phosphorylated and 9 acetylated proteins.

A distinct set of proteins were altered in response to 25(OH)D_3_ in terms of expression level and post-translational modifications. The list of altered mRNA transcripts and altered proteins following 25(OH)D_3_ treatment were aligned to determine which overlapped (**Figure 6**). Several of the genes shown to be altered at the RNA level could not be measured at the peptide level or did not have changes in protein level within this timeframe. 25(OH)D_3_ altered nine genes at both the gene expression and protein or modified protein level (p < 0.05): ***HIVEP2*** (Human Immunodeficiency Virus Type I Enhancer Binding Protein 2), a transcription factor involved in immune function and bone remodelling; ***CDRT1*** (CMT1A duplicated region transcript 1), a F-box protein that acts as protein-ubiquitin ligase; ***KIAA1522***, ***AHDC1*** (AT-Hook DNA Binding Motif Containing 1), involved in DNA binding; ***ENDOU*** (endonuclease, Poly(U) Specific) and phosphorylated ***PAK3*** (P21 (RAC1) Activated Kinase 3) a serine-threonine kinase that forms an activated complex with GTP-bound RAS-like (P21), CDC2 and RAC1 important for cytoskeletal remodelling. There was decreased gene and protein expression of ***PLEKHG2***, methylated ***HBG1*** (fetal haemoglobin) and acetylated matrix metalloproteinase-3 (***MMP3***), which is involved in the breakdown of extracellular matrix proteins, tissue remodelling and can also act as a transcription factor.

**Figure 6.**
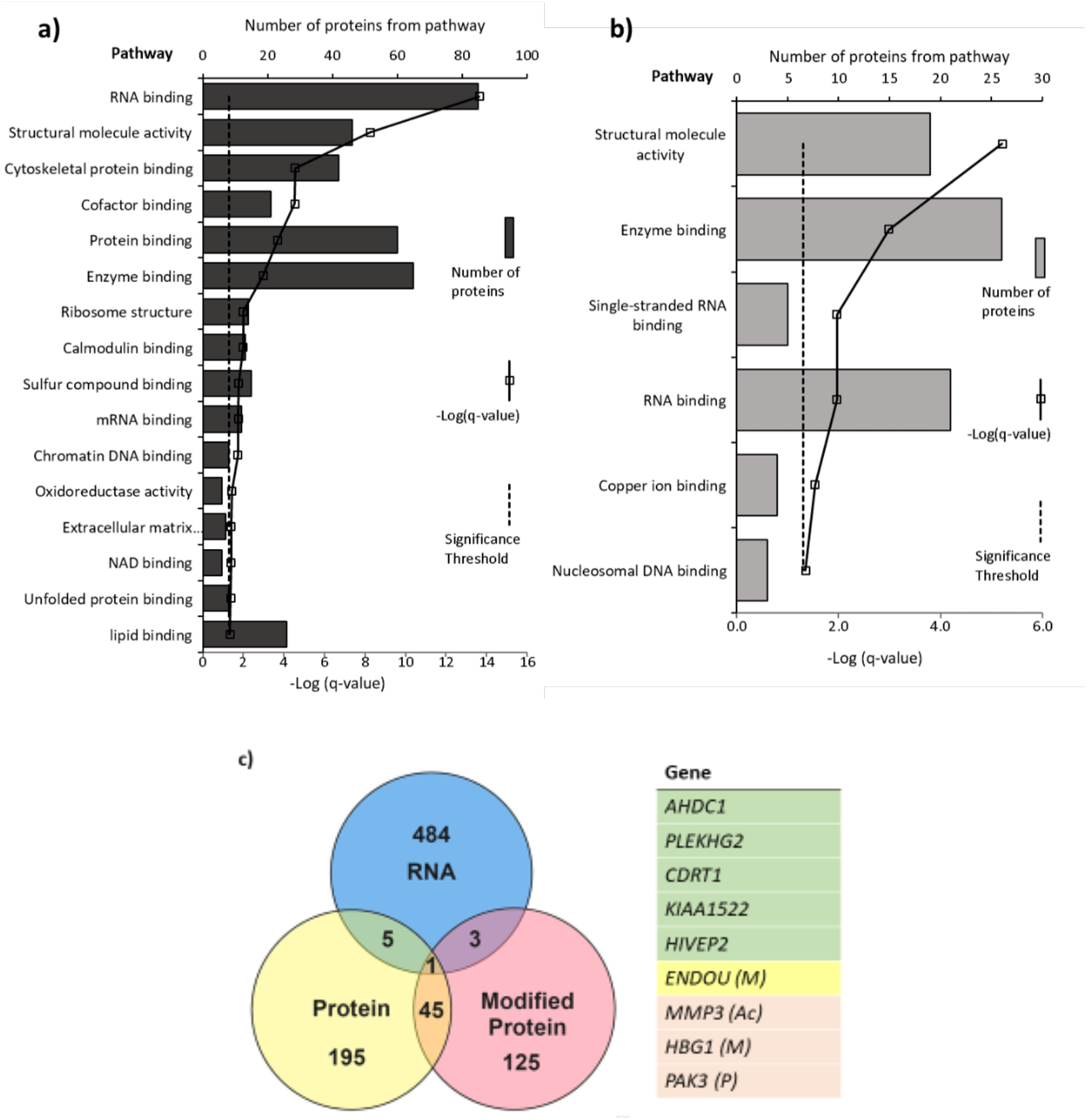
Pathways with **(a)** altered protein and **(b)** methylated protein expression in response to 25(OH)D_3_ treatment. Significantly altered pathways from genes mapped to sites of altered protein expression. Pathways identified using Toppgene and displayed as –Log q-value. **c)** Alignment of RNA and protein expression data. 9 genes were altered at both the RNA and protein level. M = methylated protein, Ac = acetylated protein, P = phosphorylated protein.

Pathways significantly enriched in the differentially expressed proteins in 25(OH)D_3_-treated placenta compared to control include: ***RNA binding, cytoskeletal protein binding and DNA chromatin binding*** (e.g. histone proteins) (FDR corrected p-value < 0.05; **Figure 6**). The methylated proteins that were differentially expressed are involved in regulating RNA, DNA and enzyme binding. One such protein group that showed altered expression was histone proteins and included methylation changes to these proteins (**Figure 6**).

The pathway analysis reveals consistent and convergent patterns of functional gene pathways modified by 25(OH)D_3_ at both the mRNA and protein level. These included gene pathways relating to structural organisation and remodelling, which could enhance placental functions in terms of general implantation and size, increasing the placenta’s potential to support fetal development. The effects of 25(OH)D_3_ also include functional gene pathways relating to the regulation of transcription via DNA binding and epigenetic modifications, supporting the importance of these genes in the placental response to vitamin D. Effects of vitamin D on the machinery that regulates wider gene expression provide a mechanism for vitamin D to have extensive effects on placental function. This supports the actions of vitamin D in terms of increased cellular growth and upregulated functions that would support increased fetal growth. Vitamin D metabolites may therefore regulate gene expression via epigenetic mechanisms or be determined by the underlying epigenetic landscape and chromatin state in a tissue-specific manner.

### Conclusion

This is the first quantitative study demonstrating 25(OH)D_3_ transfer and metabolism by the human placenta; with widespread effects on the placenta itself. We demonstrate an active 25(OH)D_3_ uptake mechanism on the maternal-facing side of the placenta and that the placenta influences the levels of 25(OH)D_3_ and its metabolites 24,25(OH)_2_D_3_ and 1,25(OH)_2_D_3_ in both the fetal and maternal circulations. Sub-optimal placental transport and metabolism of maternal 25(OH)D_3_ may therefore limit fetal supply and impede fetal development. In support of this, we see associations between placental expression levels of key vitamin D handling and metabolic genes and aspects of fetal size in our cohort study. Placental transfer of 25(OH)D_3_ metabolites may contribute to the increased maternal 1,25(OH)_2_D_3_ concentrations during gestation, which contribute to maternal physiological adaptations that support pregnancy. Effects of vitamin D on fetal development may also be mediated via effects on the placenta itself. Indeed, we show that 25(OH)D_3_ exposure differentially affects the expression of placental genes and proteins, which together map to multiple gene pathways critical for the placenta’s role in pregnancy. Epigenetic analyses suggest that the underlying epigenetic landscape helps to dictate the placental transcriptional response to vitamin D treatment, raising the possibility that environmentally induced epigenetic alterations may re-shape the vitamin D response.

These data demonstrate a complex interplay between vitamin D and the placenta to ensure that they can both play optimal roles in supporting fetal growth and maternal adaptations to pregnancy. As key mediators of this interplay, such as CYP27B1/CYP24A1/VDR, have higher expression levels earlier in pregnancy these effects will be important throughout gestation. This work therefore strongly suggests a role for placental transport and metabolism in mediating the balance of vitamin D distribution and utilisation throughout pregnancy. Understanding the regulatory mechanisms and rate-limiting steps in the relationship between vitamin D and the placenta will be a prerequisite for identifying options for targeted intervention to improve pregnancy outcomes.

## Materials and Methods

### Placental samples

For placental perfusion, placental fragment culture and primary term human cytotrophoblast culture experiments, human placentas were collected from term pregnancies immediately after delivery, following written informed consent and with the approval of the South and West Hants Local Research Ethics Committee (11/SC/0529). The study conformed to the Declaration of Helsinki.

To investigate associations between placental gene expression and maternal or fetal characteristics we used data and samples from the SWS, a cohort study of 3,158 live singleton births [28]. All data were collected with the approval of the South and West Hants Local Research Ethics Committee (276/97, 307/97). Written informed consent was obtained from all participating women and by parents or guardians with parental responsibility on behalf of their children.

### Placental perfusion

Placentas (n = 5) were perfused using the isolated perfused placental cotyledon methodology [34], as previously described [17]. Catheters (Portex, Kent, UK), 15 cm in length were inserted in the fetoplacental artery (polythene tubing: i.d. 1.0 mm, o.d. 1.6 mm) and fetoplacental vein (PVC tubing: i.d. 2 mm, o.d. 3 mm) of an intact cotyledon and sutured in place. On the maternal side, five 10 cm lengths of polythene tubing (Portex; i.d. 0.58 mm, o.d. 0.96 mm) were inserted through the decidua and into the intervillous space. The fetal circulation and intervillous space were perfused with a modified Earle’s bicarbonate buffer (EBB, 1.8 mM CaCl_2_, 0.4 mM MgSO_4_, 116.4 mM NaCl, 5.4mM KCl, 26.2 mM NaHCO_3_, 0.9 mM NaH_2_PO_4_, 5.5 mM glucose) containing 0.1% bovine serum albumin, and 5000 IU L^−1^ heparin equilibrated with 95% O_2_–5% CO_2_) using roller pumps (Watson Marlow, Falmouth, UK) at 6 and 14 ml min^−1^, respectively. Perfusion of the fetal circulation was established first, and, if fetal venous outflow was >95% of fetal arterial inflow, the maternal-side arterial perfusion with EBB was established 15 min later. Following 45 min of initial perfusion, the maternal arterial perfusion was switched to EBB containing 30 nM ^13^C-25(OH)D_3_ (Sigma Aldrich, US). Approximately 1 ml samples of fetoplacental and maternal venous outflow were collected at time points 30, 60, 90, 120, 150, 180, 200, 220, 240, 260, 280 and 300 min. All samples were snap frozen on dry ice and stored at −80°C. At the end of the experiment, the perfused cotyledon was weighed, snap frozen and stored at −80°C.

### Vitamin D metabolite measures

^13^C-25(OH)D_3_, ^13^C-24,25(OH)_2_D_3_ and ^13^C-1,25(OH)_2_D_3_ were quantified in samples using ultraperformance liquid chromatography hyphenated with electrospray ionisation – tandem mass spectrometry (UPLC-MS/MS) [35]. Analysis of extracted, placental vitamin D metabolites was performed on a Waters ACQUITY ultra performance liquid chromatography (UPLC) coupled to a Waters Xevo TQ-XS mass spectrometer [36]. 1,25(OH)_2_D_3_ in placental perfusate samples was measured using the 1,25-dihydroxy vitamin D EIA kit (Immunodiagnostic systems; UK).

### Placental fragment culture

Villous tissue fragments of approximately 10 mg were dissected from the placenta and cultured at 37°C in Tyrode’s buffer. Following the experiments as described below, buffer was removed, fragments washed in Tyrode’s buffer, snap frozen on dry ice and stored at −80°C.

#### Gene expression experiments for effects of vitamin D

Villous fragments were incubated for 8 h at 37°C in Tyrode’s buffer containing 20 μM 25(OH)D_3_ (Cayman Chemical, Michigan, USA) with 0.7 mM bovine serum albumin (BSA; Sigma Aldrich), or control buffer with 0.7 mM BSA (n = 4–15 in triplicate per treatment).

#### Gene expression experiments for vitamin D uptake

Villous fragments were incubated for 8 h at 37°C in Tyrode’s buffer containing 20 μM 25(OH)D_3_ and BSA, plus the endocytic inhibitors 5 mM amiloride hydrochloride hydrate (Sigma Aldrich), 10 μM cytochalasin D (Sigma Aldrich) or 2 μM Receptor Associated Protein (RAP; Innovative Research, USA), which block pinocytosis, classical clathrin-mediated endocytosis and megalin-mediated endocytosis, respectively (n = 4–5 in triplicate per treatment). Villous fragments had a prior 30 min pre-incubation in the specific blocker at 37°C before the 8 h incubation. Control villous fragments were incubated in Tyrode’s buffer at 37°C for the pre-incubation period and then a further 8 h.

#### DNA methylation, RNA-seq and proteomic experiments

Villous fragments were incubated for 8 h at 37°C in Tyrode’s buffer containing 20 μM 25(OH)D_3_ plus 0.7 mM BSA or control buffer with 0.7 mM BSA (n = 8 in triplicate per treatment).

#### Experiments for microscopy

Villous fragments were incubated with 150 nM FITC-albumin in Tyrode’s buffer at 4°C and 37°C. Fragments were incubated for 5, 15 or 30 min at each temperature (n = 5 per treatment group). Control fragments were also incubated with 1.43 μM FITC-dextran at 4°C and 37°C for 30 min (n = 4 at both temperatures). At each time point, buffer was removed, and villous fragments were fixed in 4% paraformaldehyde and stored at 4°C. After 24 h, fragments were transferred to 0.1% sodium azide and stored at 4°C. Fragments were further stained for villous stroma by rhodamine-PSA (*Pisum sativum* agglutinin) and the microvillous membrane of the syncytiotrophoblast by biotin-DSL (*Datura stramonium* lectin) and visualised on a SP5 fluorescent confocal microscope. Images were obtained through a series of z-sections through the fragment. The average FITC-protein uptake for each placental fragment was calculated using the software ImageJ.

### Southampton Women’s Survey (SWS) characterised placentas

Non-pregnant women aged 20–34 years were recruited to the SWS; assessments of maternal anthropometry were performed at study entry and at 11 and 34 weeks’ gestation in those who became pregnant. A tape measure was used to measure mid-upper arm circumference from which arm muscle area was derived using the formula ((mid arm circumference - π × triceps skinfold thickness)^2^/4π] - 6.5) [37]. Measures of fetal size were determined at 19 and 34 weeks’ gestation using a high-resolution ultrasound system (Acuson 128 XP, Aspen and Sequoia Mountain View, CA, USA). Standardised anatomical landmarks were used to measure head circumference, abdominal circumference, crown-rump length and femur length. Royston’s method was used to derive z-scores for ultrasound measurements of size. The method corrects for variation in age at measurement [38]. At birth, neonatal anthropometric measures were recorded (birth weight, placental weight, head circumference and crown–heel length). Within 2 weeks of birth, a subset of babies had dual energy X-ray absorptiometry (DXA) measurements of lean and fat mass using a Lunar DPX instrument with neonatal scan mode and specific paediatric software (paediatric small scan mode, v4.7c, GE Corp., Madison, WI, USA).

Placentas were collected from term SWS pregnancies within 30 minutes of delivery. Placental weight was measured after removing blood clots, cutting the umbilical cord flush with its insertion into the placenta, trimming away surrounding membranes and removing the amnion from the basal plate. Five villous tissue samples were selected from each placenta using a stratified random sampling method (to ensure that the selected samples were representative of the placenta as a whole); the maternal decidua was cut off of each sample. Samples were snap frozen in liquid nitrogen and stored at −80°C. For this study, a cohort of 102 placentae was selected from 300 collected in total based on availability of neonatal data. For each placenta, the five samples were pooled and powdered in a frozen tissue press.

### RNA and DNA extraction

RNA was extracted from 30 mg placental samples using the miRNeasy mini kit with the RNase-free DNase Set (Qiagen, UK) according to manufacturer’s instructions. DNA was extracted from placental fragments using the Qiagen DNeasy Blood & Tissue Kit (Qiagen, UK) according to manufacturer’s instructions. RNA and DNA were quantified by UV absorption (NanoDrop 1000, Thermo Scientific, UK). RNA quality was assessed with an RNA 2100 Bioanalyser (Agilent, USA) and accepted if the RNA integrity number (RIN) was above 6.0.

### Quantitative reverse transcription PCR (qRT-PCR)

Total RNA (0.2 μg) was reverse transcribed into cDNA and gene expression was measured using qRT-PCR with a Roche Light-Cycler-480. Oligonucleotide probes were supplied by Roche (Universal Probe Library (UPL); Roche, UK) and primers supplied by Eurogentec (Seraing, Belgium). Control genes were selected using the geNorm™ human Housekeeping Gene Selection Kit (Primer Design Limited, Southampton UK). For UPL probes cycle parameters were 95°C for 10 min; 45 cycles of 95°C for 10 s, 60°C for 30 s; then 72°C for 1 s. For Perfect Probes, the cycle parameters were 95°C for 10 min; 50 cycles of 95°C for 15 s, 60°C for 30 s and 72°C for 15 s. Intra-assay coefficients of variation for each gene were 5–8%. All samples were run on the same plate in triplicate. All SWS placenta mRNA levels are presented relative to the geometric mean of the three control genes, tyrosine 3-monooxygenase/tryptophan 5-monooxygenase activation protein, zeta polypeptide (*YWHAZ*), ubiquitin C (*UBC*) and topoisomerase (*TOP1*) [39]. Placental fragment gene expression data was normalised to the geometric mean of *YWHAZ* and *UBC* as *TOP1* expression was altered by vitamin D treatment.

### RNA sequencing

Placental fragment RNA samples (450 ng) were converted into cDNA libraries using the Illumina TruSeq Stranded mRNA sample preparation kit. Stranded RNA sequencing was carried out by Expression Analysis (Durham, USA) using HiSeq 2×50bp paired-end sequencing on an Illumina platform. After quality control, reads were mapped to hg38 (Ensembl; March 2017) using HISAT2 v2.0.5 [40] and counted with HTSeq v0.6.1p1 [41] using union mode and stranded=reversed settings. Genes were filtered based upon counts per million (CPM) and genes with < 1 CPM in at least half of the samples were removed. Samples were normalised using the trimmed mean of M-values (TMM) method, and median plots demonstrated no samples outside two standard deviations. Differential expression analysis was carried out using EdgeR [42] in the R statistical computing environment (https://www.R-project.org/). Data were filtered to a Benjamini & Hochberg (B&H) false discovery rate (FDR) corrected probability of 5%. Genes with altered expression were mapped to pathways using Toppgene (Division of Bioinformatics, Cincinnati Children’s Hospital Medical Centre) [43]. Pathways were accepted with a minimum hit count of 4 and B&H FDR corrected q value of ≤ 0.05.

### Illumina 850K DNA methylation

DNA methylation analysis was carried out using the Infinium^®^ MethylationEPIC array at Barts and the London School of Medicine and Dentistry Genome Centre. Methylation data as β-values were normalised and differentially methylated CpGs were identified by a Wilcoxon signed-rank test with control versus vitamin D treated, using a 10% change cut-off. Sites were removed that contain any missing values. All samples met minimal inclusion criteria for analysis, as each sample had >75% sites with a detection p value <1×10^−5^. Probes on X and Y chromosomes were removed. The average CpG methylation levels were calculated for all gene promoters (TSS ± 2 kb).

RNA sequencing data, ChIP-seq data and methylation array data that support the findings of this study have been deposited in the Gene Expression Omnibus (GEO) with the accession code GSE167431.

### Proteomics

#### Quantitative proteomics sample processing

Placental villous fragments were snap frozen at −80 °C. These were dissolved in 0.5 M triethylammonium bicarbonate, 0.05% sodium dodecyl sulphate and subjected to pulsed probe sonication (Misonix, Farmingdale, NY, USA). Lysates were centrifuged (16,000 g, 10 min, 4°C) and supernatants measured for protein content using infrared spectroscopy (Merck Millipore, Darmstadt, Germany). Lysates were then reduced, alkylated and subjected to trypsin proteolysis. Peptides were isobaric stable isotope labelled using the eight-plex iTRAQ reagent kit and analysed using orthogonal 2-dimensional liquid chromatography (offline alkaline C4 reverse phase and online acidic C18 reverse phase) hyphenated with nanospray ionisation and high-resolution tandem mass spectrometry, as previously reported by the authors [44, 45].

#### Database searching

Unprocessed raw files were submitted to Proteome Discoverer 1.4 (Thermo Fisher Scientific) for target decoy searching against the UniProtKB homo sapiens database (https://www.uniprot.org/proteomes/UP000005640), which comprised of 20,159 entries (release date January 2015), allowing for up to two missed cleavages, a precursor mass tolerance of 10 ppm, a minimum peptide length of 6 and a maximum of two variable (1 equal) modifications of 8-plex iTraq (Y), oxidation (M), deamidation (N, Q), or phosphorylation (S, T, Y). Methylthio (C) and iTraq (K, Y and N-terminus) were set as fixed modifications. FDR at the peptide level was set at < 0.05. Percent co-isolation excluding peptides from quantitation was set at 50. Reporter ion ratios from unique peptides only were taken into consideration for the quantitation of the respective protein. A permutation test using the normalised iTraq ratios of each vitamin D-treated placenta compared to its respective control was performed. Significance was set at p < 0.05. In adherence to the Paris Publication Guidelines for the analysis and documentation of peptide and protein identifications (https://doi.org/10.1074/mcp.T400006-MCP200), only proteins identified with at least 2 unique peptides were further subjected to bioinformatics. All mass spectrometry data have been deposited to the ProteomeXchange Consortium via the Proteomics Identifications Archive with the data set identifier PXD011443. Proteins with altered expression were mapped to gene pathways using Toppgene (Division of Bioinformatics, Cincinnati Children’s Hospital Medical Centre). Pathways were accepted with a minimum hit count of 4 and FDR corrected q value of ≤ 0.05.

### Primary term human cytotrophoblast culture

Cytotrophoblast cells were isolated from term human placentas (n = 2) using an adaptation of the method developed by Kliman *et al*. (1986) [46], as described previously [32]. Isolated cells were plated in culture medium (Dulbecco’s modified Eagle’s medium and Ham’s F-12 1:1, 10% heat inactivated fetal calf serum, 0.6% glutamine and the antibiotics 1% gentamicin, 0.2% penicillin and 0.2% streptomycin) onto 35 mm culture dishes (Nunc), at a density of 2.5 × 10^6^, and were maintained in primary culture at 37°C in a humidified incubator (95% air–5% CO_2_). At 66 h, matched cell samples had control media or 25(OH)D_3_ added to a final concentration of 20 μM and were cultured for a period of 24 h before collection at 90 h. β-hCG released by cultured cytotrophoblast cells into the culture media was measured as a marker of syncytialisation using a β-hCG ELISA Kit. Increasing levels of β-hCG production were observed by 66 h of culture; average β-hCG levels in the 90 h culture media were 413.13 mIU/ml. At 90 h media was removed, cells were washed in PBS and fixed in methanol. Fixed cytotrophoblast cells were blocked with BSA and incubated overnight at 4°C with 1/100 mouse anti-human Desmoplakin antibody. Cells were washed and incubated for 2 h at RT with 1/250 DAPI + 1/500 goat anti-mouse Alexa Fluor 568. Cells were again washed and stored under PBS at 4°C until imaging on a Lecia SP8 confocal microscope.

### ChIP-seq

Control and Vitamin D treated cytotrophoblast cells were fixed with 1% formaldehyde for 12 min in PBS. After quenching with glycine (final concentration 0.125 M), fixed cells were washed and lysed as previously described (Latos et al., 2015). Chromatin was sonicated using a Bioruptor Pico (Diagenode), to an average size of 200–700 bp. Immunoprecipitation was performed using 10 μg of chromatin and 2.5 μg of antibody (H3K4me3 (Diagenode C15410003), H3K4me1 (abcam ab8895 and H3K27ac (Diagenode C15410196). Final DNA purification was performed using the GeneJET PCR Purification Kit (Thermo Scientific, K0701) and eluted in 80 μL of elution buffer. ChIP-seq libraries were prepared from 1 to 5 ng eluted DNA using NEBNext Ultra II DNA library Prep Kit (New England BioLabs), with 12 cycles of library amplification. Libraries were sequenced on an Illumina NextSeq 500 with single-ended 75 bp reads at the Barts and the London Genome Centre. Sequencing reads were trimmed with Trim Galore, mapped to hg38 genome assembly using bowtie2, and filtered to retain uniquely mapped reads. Peak detection was performed using MACS2 with the --broad option. Using DiffBind, we detected no peaks with significant differences in enrichment of either histone mark upon vitamin D treatment. ChIP-seq signal at gene promoters (±1kb from TSS) was measured as reads per million.

### External data

ENCODE ChIP-seq data from 16-week placentas were extracted as processed peaks (ENCFF707YUS, ENCFF180ADH). RNA-seq data from THP1 cells (GSE69284) were extracted as processed differentially expressed genes. Raw sequencing reads from FAIRE-seq data in THP1 cells (GSE69297) were downloaded and processed as for ChIP-seq data.

### Data analysis

Data are presented as mean (SEM) and analysed using IBM^®^ SPSS^®^ Statistics Version 20 (IBM^®^, Armonk, New York, USA), unless otherwise stated. Data were log transformed if not normally distributed. Significance was set at p < 0.05. A two-way ANOVA was used to explore the effects of temperature and time on FITC-albumin uptake in the fragment experiments. qRT-PCR data were analysed with one-way ANOVA and Tukey’s post-hoc test.

For the SWS data, maternal variables that were not normally distributed were transformed logarithmically.

Partial correlations of placental gene expression levels and maternal, fetal or neonatal variables were calculated, controlling for sex. Neonatal variables were additionally adjusted for gestational age.

Differences in epigenomic data (EPIC arrays, ChIP-seq, FAIRE-seq) at gene promoters were evaluated using nonparametric Wilcoxon tests, with Benjamini-Hochberg correction for multiple comparisons.

## Acknowledgments

CS was funded by a Gerald Kerkut Charitable Trust studentship and BA by Rank Prize and University of Southampton Vice Chancellor’s Studentships plus the MRC.

KMG is supported by the UK Medical Research Council (MC_UU_12011/4), the National Institute for Health Research (NIHR Senior Investigator (NF-SI-0515-10042), NIHR Southampton 1000DaysPlus Global Nutrition Research Group (17/63/154) and NIHR Southampton Biomedical Research Centre (IS-BRC-1215-20004)), British Heart Foundation (RG/15/17/3174) and the US National Institute On Aging of the National Institutes of Health (Award No. U24AG047867).

KSJ is supported by the National Institute for Health Research (NIHR) Cambridge Biomedical Research Centre (IS-BRC-1215-20014). The NIHR Cambridge Biomedical Research Centre is a partnership between Cambridge University Hospitals NHS Foundation Trust and the University of Cambridge, funded by the NIHR. The views expressed are those of the authors and not necessarily those of the NHS, the NIHR or the Department of Health and Social Care. Experimental work performed by KSJ and FH at MRC EWL was supported by Dr Ann Prentice (UK Medical Research Council U105960371).

The SWS has been supported by grants from Medical Research Council (MRC) [4050502589 (MRC LEU)], Dunhill Medical Trust, British Heart Foundation, Food Standards Agency, National Institute for Health Research (NIHR) Southampton Biomedical Research Centre, University of Southampton and University Hospital Southampton NHS Foundation Trust, NIHR Oxford Biomedical Research Centre, University of Oxford and the European Union’s Seventh Framework Programme (FP7/2007-2013), project EarlyNutrition, under grant agreement 289346 and the European Union’s Horizon 2020 research and innovation programme (LIFECYCLE, grant agreement No 733206).

EC has been supported by the Wellcome Trust (201268/Z/16/Z). Work leading to these results was supported by the BBSRC (HDHL-Biomarkers, BB/P028179/1), as part of the ALPHABET project, supported by an award made through the ERA-Net on Biomarkers for Nutrition and Health (ERA HDHL), Horizon 2020 grant agreement number 696295.

The proteomic analyses (SDG and AM) were financially supported by the National Institutes of Health (R21AI122389) and the Beckman Institute at the California Institute of Technology. We acknowledge funding from the European Union H2020-MSCA-IF-2018 to JMF under grant agreement no. 841172 (InvADeRS).

The electron microscopy image in Figure 2 was produced with help of the Biomedical imaging unit, Faculty of Medicine, University of Southampton. EC has been supported by the Wellcome Trust (201268/Z/16/Z) and an NIHR Clinical Lectureship.

