## supporting information for "Placental uptake and metabolism of 25(OH)Vitamin D determines its activity within the fetoplacental unit"

### Placental uptake and metabolism as determinants of pregnancy vitamin D status

#### SUPPORTING INFORMATION

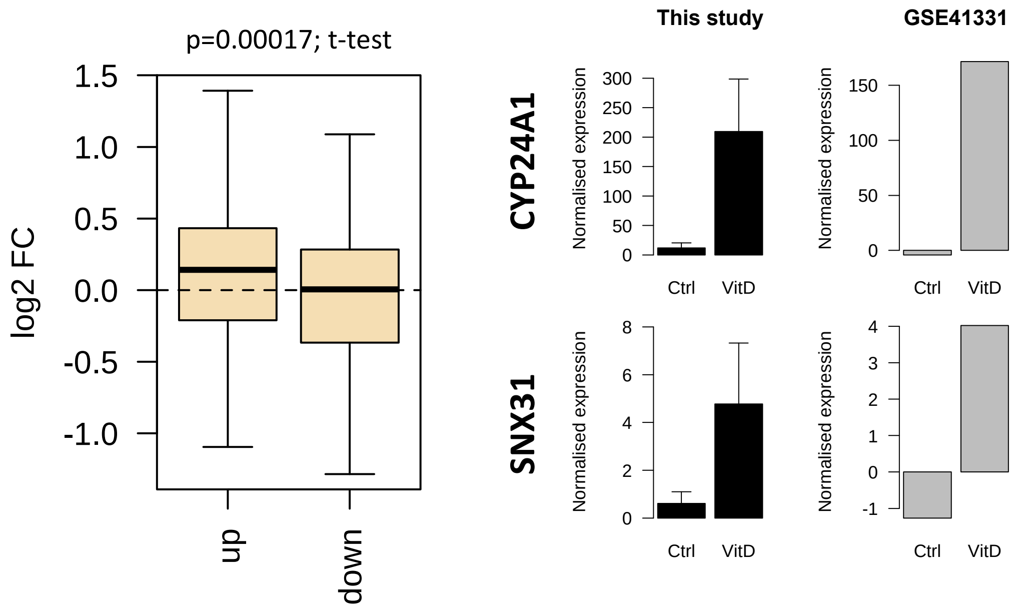


**Supplementary Figure 1.** Gene expression changes observed in this study following 8 h 25(OH)$D_3$ exposure were compared with those in a published dataset from human placenta (GSE41331), which looked at longer-term vitamin D response (24 h). Genes that were upregulated following 8 h exposure also tended to have increased expression in the 24 h dataset, whereas downregulated genes were unchanged. Differentially expressed genes in common between both placental datasets included *CYP24A1* and *SNX31*.

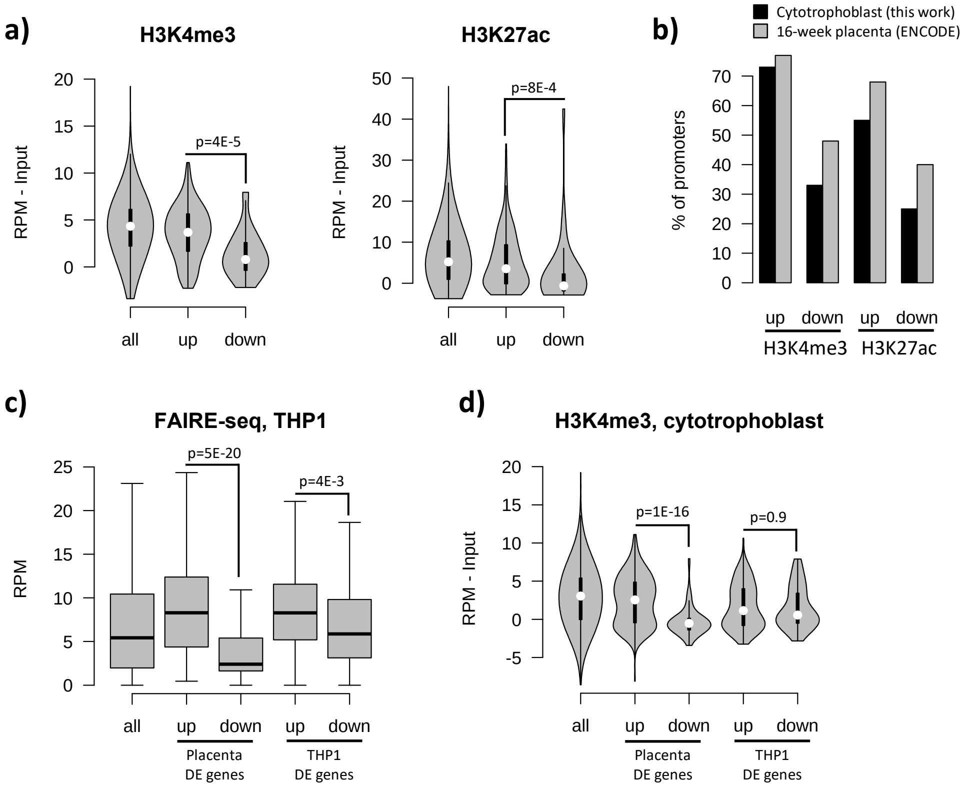


**Supplementary Figure 2. a)** Promoters of upregulated genes displayed higher levels of both H3K4me3 and H3K27ac than those seen at downregulated genes: this pattern was also seen when focusing only on CpG island promoters. **b)** Although our ChIP-seq data is from isolated cytotrophoblast, very similar patterns were observed in ENCODE data from 16-week placenta. **c)** Open chromatin (FAIRE-seq) data from THP1 cells treated with $1,25(OH)_2D_3$ for 4 h. The promoters of both placenta- and THP1-upregulated genes displayed open chromatin in THP1 cells. **d)** H3K4me3 levels in the placenta were higher for promoters of placenta-upregulated genes than for THP1-upregulated ones.

**Supplementary Table 1. RNA-seq derived gene signatures of human placental samples following 8 h 25(OH)$D_3$ incubation.** 493 genes were differentially expressed between 25(OH)$D_3$ treated compared to control placental fragments (358 increased and 135 decreased) at a FDR adjusted p value < 0.05.

| Genes | Symbol | logFC | FDR |
|---|---|---|---|
| ENSG00000019186 | *cyp24a1* | 4.406502 | 1.07E-31 |
| ENSG00000123612 | *ACVR1C* | 3.317085 | 1.08E-17 |
| ENSG00000135436 | *FAM186B* | 3.016757 | 1.71E-14 |
| ENSG00000174226 | *SNX31* | 3.01061 | 3.99E-17 |
| ENSG00000179564 | *LSMEM2* | 2.549959 | 3.72E-09 |
| ENSG00000250519 | | 2.544841 | 5.08E-08 |
| ENSG00000105371 | *ICAM4* | 2.331246 | 6.46E-13 |
| ENSG00000255202 | *LOC10537661* | 2.248171 | 3.49E-07 |
| ENSG00000215483 | *LINC00548* | 2.238101 | 3.89E-11 |
| ENSG00000280054 | | 2.034102 | 2.61E-06 |
| ENSG00000273599 | | 2.027395 | 1.5E-06 |
| ENSG00000164404 | *GDF9* | 1.997981 | 1.19E-06 |
| ENSG00000008300 | *MIR4793* | 1.983139 | 8.71E-06 |
| ENSG00000185900 | *POMK* | 1.925198 | 1.85E-08 |
| ENSG00000230069 | | 1.909871 | 1.86E-05 |
| ENSG00000228037 | *LOC10099658* | 1.897453 | 5.28E-06 |
| ENSG00000178502 | *KLHL11* | 1.832096 | 6.31E-11 |
| ENSG00000243629 | *LINC00880* | 1.828392 | 5.37E-05 |
| ENSG00000240032 | | 1.826124 | 3.41E-06 |
| ENSG00000235499 | | 1.784459 | 0.001338 |
| ENSG00000237927 | | 1.772789 | 0.000109 |
| ENSG00000105376 | *ICAM5* | 1.75958 | 5.82E-10 |
| ENSG00000231966 | | 1.723246 | 0.000759 |
| ENSG00000232586 | | 1.702729 | 1.67E-05 |
| ENSG00000160051 | *IQCC* | 1.699603 | 5.47E-07 |
| ENSG00000188167 | *TMPPE* | 1.674347 | 1.54E-06 |
| ENSG00000279283 | | 1.672608 | 0.000262 |
| ENSG00000186472 | *PCLO* | 1.658666 | 0.000331 |
| ENSG00000246560 | *LOC10537734* | 1.656954 | 0.000196 |
| ENSG00000165194 | *PCDH19* | 1.646677 | 8.45E-06 |
| ENSG00000233930 | *KRTAP5-AS1* | 1.632016 | 0.000777 |
| [illegible] | | [illegible] | [illegible] |

DE genes UP | DE genes DOWN

**Supplementary Table 2. DNA methylation changes in human placental samples following 8 h 25(OH)$D_3$ incubation.** 319 CpGs displaying methylation differences larger than 10% (230 hypomethylated, 89 hypermethylated).

| Chromosome | Start | End | Feature | Feature Strand | Feature Orientation | Distance |
|---|---|---|---|---|---|---|
| 1 | 2707574 | 2707574 | TTC34 | - | downstream | 294 |
| 1 | 2767082 | 2767082 | | | Not found | 0 |
| 1 | 5976041 | 5976041 | | | Not found | 0 |
| 1 | 7018950 | 7018950 | | | Not found | 0 |
| 1 | 7120927 | 7120927 | | | Not found | 0 |
| 1 | 7181418 | 7181418 | | | Not found | 0 |
| 1 | 16347985 | 16347985 | CLCNKA | + | overlapping | 0 |
| 1 | 91301732 | 91301732 | RP4-665J23.1 | - | overlapping | 0 |
| 1 | 96124811 | 96124811 | | | Not found | 0 |
| 1 | 175048874 | 175048874 | TNN | + | upstream | 1349 |
| 1 | 194458735 | 194458735 | | | Not found | 0 |
| 1 | 208900752 | 208900752 | RP11-459K23.2 | + | downstream | 281 |
| 2 | 817734 | 817734 | | | Not found | 0 |
| 2 | 1498139 | 1498139 | | | Not found | 0 |
| 2 | 1748618 | 1748618 | PXDN | - | overlapping | 0 |
| 2 | 1932997 | 1932997 | | | Not found | 0 |
| 2 | 2337094 | 2337094 | MYT1L | - | downstream | 1062 |
| 2 | 2875087 | 2875087 | AC011995.3 | - | overlapping | 0 |
| 2 | 27938293 | 27938293 | AC074091.13 | - | overlapping | 0 |
| 2 | 27938372 | 27938372 | AC074091.13 | - | overlapping | 0 |
| 2 | 27938765 | 27938765 | AC074091.13 | - | overlapping | 0 |
| 2 | 34628357 | 34628357 | | | Not found | 0 |
| 2 | 79878369 | 79878369 | CTNNA2 | + | overlapping | 0 |
| 2 | 126468026 | 126468026 | AC097499.1 | + | overlapping | 0 |
| 2 | 144355866 | 144355866 | RP11-570L15.1 | - | overlapping | 0 |
| 2 | 145606432 | 145606432 | | | Not found | 0 |
| 2 | 147345338 | 147345338 | | | Not found | 0 |
| 2 | 177028622 | 177028622 | HOXD3 | + | overlapping | 0 |
| 2 | 181370099 | 181370099 | | | Not found | 0 |
| 2 | 199240044 | 199240044 | AC005235.1 | - | overlapping | 0 |
| 2 | 202982522 | 202982522 | AC079354.3 | - | overlapping | 0 |
| 2 | 217675070 | 217675070 | | | Not found | 0 |
| 3 | 88031396 | 88031396 | | | Not found | 0 |

hypermethylation | hypomethylation
